# Modeling cell size regulation under complex and dynamic environments

**DOI:** 10.1101/2022.09.09.507356

**Authors:** César Nieto, César Vargas-García, Juan Manuel Pedraza, Abhyudai Singh

**Affiliations:** Department of Electrical and Computer Engineering,University of Delaware, Newark, DE, USA; AGROSAVIA. Corporación Colombiana de Investigación Agropecuaria, Bogotá, Colombia; Department of physics. Universidad de los Andes. Bogotá.Colombia

## Abstract

In nature, cells face changes in environmental conditions that can modify their growth rate. In these dynamic environments, recent experiments found changes in cell size regulation. Currently, there are few clues about the origin of these cell size changes. In this work, we model cell division as a stochastic process that occurs at a rate proportional to the size. We propose that this rate is zero if the cell is smaller than a minimum size. We show how this model predicts some of the properties found in cell size regulation. For example, among our predictions, we found that the mean cell size is an exponential function of the growth rate under steady conditions. We predict that cells become smaller and the way the division strategy changes during dynamic nutrient depletion. Finally, we use the model to predict cell regulation in an arbitrary complex dynamic environment.

## INTRODUCTION

In natural environments, cells face different complex and dynamic changes (Koch, 1971). Some of these dynamics can be associated with external events such as the arrival of new nutrients, fluctuations in temperature (Saarinen et al., 2018), or the competence against other species (Grover et al., 1997). A typical alteration of the environment present in all proliferating populations is the depletion of nutrients resulting from population growth (Morita, 1990).

The recent development of precise high-throughput experiments has revealed that many cell properties can change dynamically as the environment runs out of nutrients (Lennon and Jones, 2011; Bakshi et al., 2021). Among the cell properties studied in these experiments, we can highlight both the regulation of cell size and the response to stress (Bakshi et al., 2021; Shimaya et al., 2021). Although they have a detailed data set, these experiments provide little insight into the mechanisms of cell size regulation in complex environmental dynamics, mainly because models that explain these findings are scarce.

Cell size regulation can be modeled as a continuous-time stochastic hybrid system (Ghusinga et al., 2016; Vargas-Garcia et al., 2016). Its dynamics consists of a continuous growth of the cell size (Fig. 1A), alternating with stochastic jumps representing the cell division. The strategy defining cell division is traditionally studied by correlation between the size after the most recent division (size at birth) and the size just before the next division (size at division) (Amir, 2014; Taheri-Araghi et al., 2015; Sauls et al., 2016). From this correlation, division strategies for exponentially growing cells can be classified into three basic paradigms (Fig. 1B) (Sauls et al., 2016): the *sizer* strategy, where the size at division is independent of the size at birth ; the *timer*, where the size at division is highly correlated with the size at birth; and the *adder*, where the size at division is the size at birth plus a random variable (the added size) with mean independent of the size at birth (Xia et al., 2020; Amir, 2014).

**Fig. 1.**
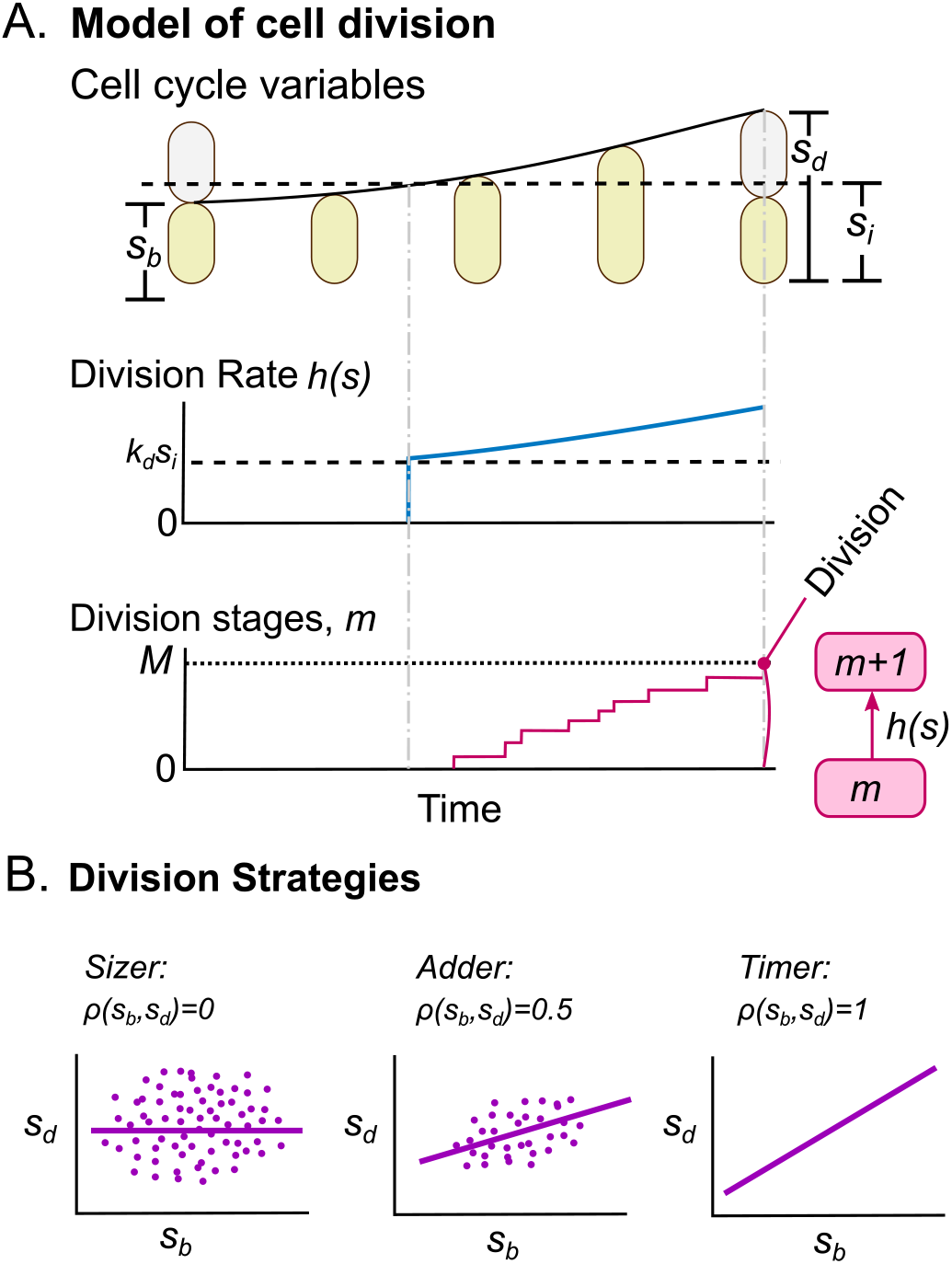
Division mechanism in cells. A. (Top) We show the main variables of the cell cycle, such as the size at birth (*s*_*b*_), the size at division (*s*_*d*_), and the minimum size to division *s*_*i*_. (center) When the cell size *s* is less than *s*_*i*_, the division rate *h*(*s*) is zero. If *s > s*_*i*_, the total division rate is *k*_*d*_*s*, with *k*_*d*_ constant. (Bottom) Division consists of a process in which cells must complete *M* division stages. The transition rate between stage *m* and stage *m* + 1 is *h*(*s*). B. Paradigms of cell division and their classification using the correlation coefficient *ρ*(*s*_*b*_, *s*_*d*_). (left) Sizer, where *s*_*d*_ is independent of *s*_*b*_. (center) Adder, where *s*_*d*_ = *s*_*b*_ + Δ with Δ being an independent random variable. (Right) Timer, where *s*_*d*_ = 2*s*_*b*_.

Current cell size regulation models have focused mainly on steady growth conditions (Wang et al., 2010; Taheri-Araghi et al., 2015; Osella et al., 2014; Amir, 2014; Chien et al., 2012; Iyer-Biswas et al., 2014a; Vargas-Garcia et al., 2018; Sanders et al., 2022; Vahdat et al., 2021). A non-steady condition of interest in cell biology is when nutrients are depleted as a result of population growth (Zwietering et al., 1990). This process is summarized as follows. When cells are inoculated into fresh medium, they grow and divide roughly in a balanced way, maintaining size stability. As a result, the biomass of the population grows exponentially. This stage is also called *exponential phase*. As nutrients begin to deplete, the growth rate decreases and the population enters the known as *stationary phase*. In this article, we model the cell size dynamics in this nutrient depletion dynamics and other more general ones.

To describe these dynamics, we model the cell size regulation as a stochastic hybrid process with continuous exponential growth and discrete divisions modeled as jumps occurring at a rate proportional to the size. Although this idea was discussed in previous studies (Jia et al., 2021; Nieto et al., 2020a; Ghusinga et al., 2016), here we include a fundamental modification: we propose that division cannot begin if the cell size is smaller than a certain minimum size. In Section 1. we present the methods; in Section 2.1, we observe how both the cell size and the division strategy depend on the growth rate for steady growth. In Section 2.2, we use this division mechanism to model cell size dynamics as nutrients are depleted, predicting results similar to those observed in experiments such as (Bakshi et al., 2021; Shimaya et al., 2021). In Section 2.3, we discuss the dynamics of both cell size and division strategy under arbitrary growth fluctuations. Section 3 presents a discussion of the limitations and interpretation of the results.

## 1. METHODS

### 1.1 The division strategy

Mathematically, (Fig. 1A) cell division mechanisms can be classified by the relationship between the size at division *s*_*d*_ and the size at birth *s*_*b*_ (Amir, 2014; Sauls et al., 2016). In the range where this relationship is linear, we can assume the following mapping:

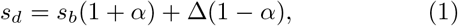

where Δ is an independent and identically distributed random variable with mean ⟨Δ⟩ *>* 0. Depending on the value of *α*, we can define the division strategy for the sizer *α* = − 1, for the adder *α* = 0, and for the timer *α* = − 1 (Campos et al., 2014; Amir, 2014).

We assume that the cells reached stability; this means ⟨*s*_*d*_⟩ = 2 ⟨*s*_*b*_⟩ with ⟨.⟩ being the expected value. From this we can derive the main properties of the division strategy. For instance, the statistics of the division control variables:

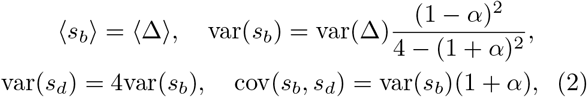

where var(*x*) = ⟨*x*^2^⟩ − ⟨*x*⟩^2^ and cov(*x, y*) = ⟨*xy*⟩ − ⟨*x*⟩ ⟨*y*⟩. From (2), we can obtain the correlation coefficient between *s*_*b*_ and *s*_*d*_:

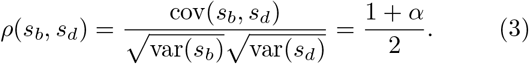

For a shorter notation, *ρ* = *ρ*(*s*_*b*_, *s*_*d*_). Fig. 1B shows that, for the sizer *ρ* = 0, while for the adder *ρ* = 0.5. Finally, for the timer *ρ* = 1. This perfect correlation for the timer strategy was previously discussed (Vargas-Garcia et al., 2016) as a condition of stability.

In *E. coli* and *B. subtilis* bacteria, under balanced and fast growth conditions, division follows the adder strategy (Taheri-Araghi et al., 2015), while under slow growth conditions, experiments show that the division strategy is sizer-like; this is 0 *< ρ <* 0.5 (Si et al., 2019; Nieto et al., 2020a). Regarding the regulation of divisions during nutrient depletion, recently it was found that the first division after leaving the stationary phase is almost sizer *ρ* ≈ 0 and subsequent divisions follow the adder *ρ* ≈ 0.5. Finally, entering the stationary phase, the strategy is close to the timer (0.5 *< ρ <* 1) (Bakshi et al., 2021).

### 1.2 Continuous-time model of cell division

This section aims to introduce a general approach for estimating the dynamics of cell size under arbitrary changing conditions. In general, cell size *s* grows at a time-dependent growth rate *µ*(*t*) according to the following:

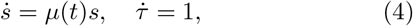

with *τ* defined as a timer that resets to zero just after each division. Table 1 presents these variables and all the others studied in this article.

**Table 1.** Variables used throughout the article.

| Variable | Interpretation |
| --- | --- |
| $s$ | Cell size. |
| $s_b$ | Cell size at the beginning of one cycle. |
| $s_d$ | Cell size at division. |
| $\mu$ | Growth rate. |
| $\mu_0$ | Reference growth rate |
| $s_i$ | Minimum size for starting the division. |
| $k_d$ | Division rate constant. |
| $M$ | Total division steps before division. |
| $m$ | Partial division steps. |
| $N$ | Nutrients in the growth medium. |
| $B$ | Total biomass of a cell population. |
| $\rho$ | Correlation coefficient between $s_b$ and $s_d$. |

If any previous division occurred at a time *t*_0_, the size *s*(*τ*) after a time *τ* since the last division follows.

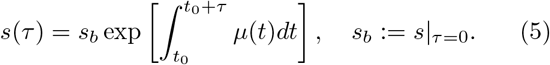

As explained in Fig. 1A, we define division as a stochastic jump process triggered after the occurrence of *M* ∈ ℕ division stages. Transitions between stages occur at a rate *h*(*s*(*τ*)) with *h*, generally a function that depends implicitly on the time through size *s*.

If *P*_*m*_(*τ*) is the probability of having completed the division stages *m* ≤ *M* at time *τ*, the division is described by the master equation:

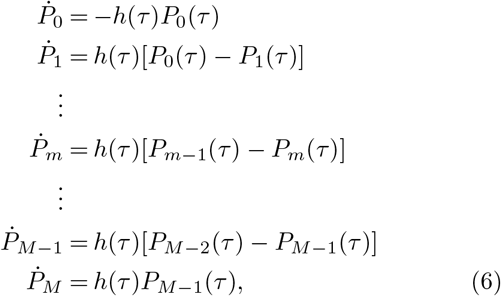

with initial conditions 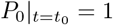 and 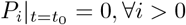.

This means that, after each division, cells reset their division stages to *m* = 0.

The probability density function (PDF) 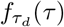 that describes the time-to-divide distribution *τ*_*d*_ follows:

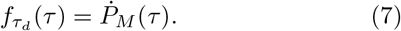

After obtaining 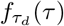, we resort to simulation methods similar to the one explained in (Thomas and Shahrezaei, 2021; Nieto et al., 2021; Blanco et al., 2020). For each cycle, we generate a random *τ*_*d*_ distributed as (7). After defining *τ*_*d*_, we estimate the dynamics of cell size throughout the cell cycle by integration of expression (5). We do this process for all the simulated cells during the total simulation time.

We consider that the cell does not split exactly in half (Iyer-Biswas et al., 2014b; Modi et al., 2017). In our simulations, we implement this property by multiplying the size by a random variable with mean 0.5 and distributed as a beta distribution with standard deviation *σ*_*div*_ = 0.05. After division, we track only one of the descendant cells, discarding the other one. With this, the cell population is constant.

### 1.3 The Minimum Size for starting division

Recent experimental findings conclude that chromosome replication and division can begin only after a minimum size is reached (Ho and Amir, 2015; Campos et al., 2014; Wallden et al., 2016; Si et al., 2017). Based on these studies, we propose in Fig. 1B, that the division rate *h*, given in (6), is a size-dependent function following:

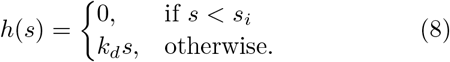

We can compare this model with a traditional division strategy, the adder, as has been studied in previous articles (Ghusinga et al., 2016; Nieto et al., 2021). To obtain the adder, we consider just *h*(*s*) = *k*_*d*_*s* with no minimum size. The PDF *f* (*s*_*d*_|*s*_*b*_) of *s*_*d*_ given *s*_*b*_ follows (Nieto et al., 2020a):

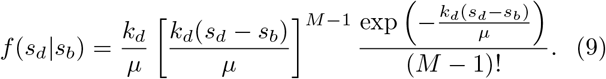

This distribution depends only on the difference Δ = *s*_*d*_ − *s*_*b*_. Therefore, by defining 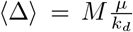, we observe that the mean size at division, given the size at birth, follows the formula:

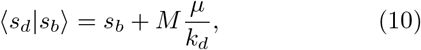

which is interpreted as an adder division strategy since ⟨*s*_*d*_⟩ is just *s*_*b*_ plus a constant.

If we consider the steady case where ⟨*s*_*d*_⟩ = 2 ⟨*s*_*b*_⟩, we find that

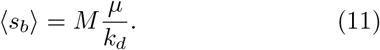

After solving ⟨*s*_*b*_⟩, we observe that our model depicted in (8) is equivalent to the adder in the limit *s*_*i*_ ≪ ⟨*s*_*b*_⟩, or more explicitly 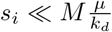. This means that the adder strategy emerges if the cell grows at a rate *µ* much faster than the division constant *k*_*d*_*/M*. In the following sections, we will observe how the division strategy deviates from the adder as the growth rate changes relatively to *k*_*d*_, especially in cases where ⟨*s*_*b*_⟩ ≈ *s*_*i*_.

## 2. RESULTS

### 2.1 Size regulation in steady growth conditions

In the first part of our study, we observe how the division regulation changes with the growth rate under steady conditions. To do this, we use the minimum size division model *s*_*i*_ considering different stable growth rates *µ*(*t*) = *µ* while keeping the division constant *k*_*d*_.

Fig. 2A shows how the main variables that quantify size statistics: the mean size at birth ⟨*s*_*b*_⟩, the size at division ⟨*s*_*d*_⟩ and the mean size ⟨*s*⟩ depend on the growth rate. As a main result, we can observe that these sizes follow an exponential function of the growth rate, as observed by most experiments (Ho and Amir, 2015; Schaechter et al., 1958; Soifer et al., 2016).

**Fig. 2.**
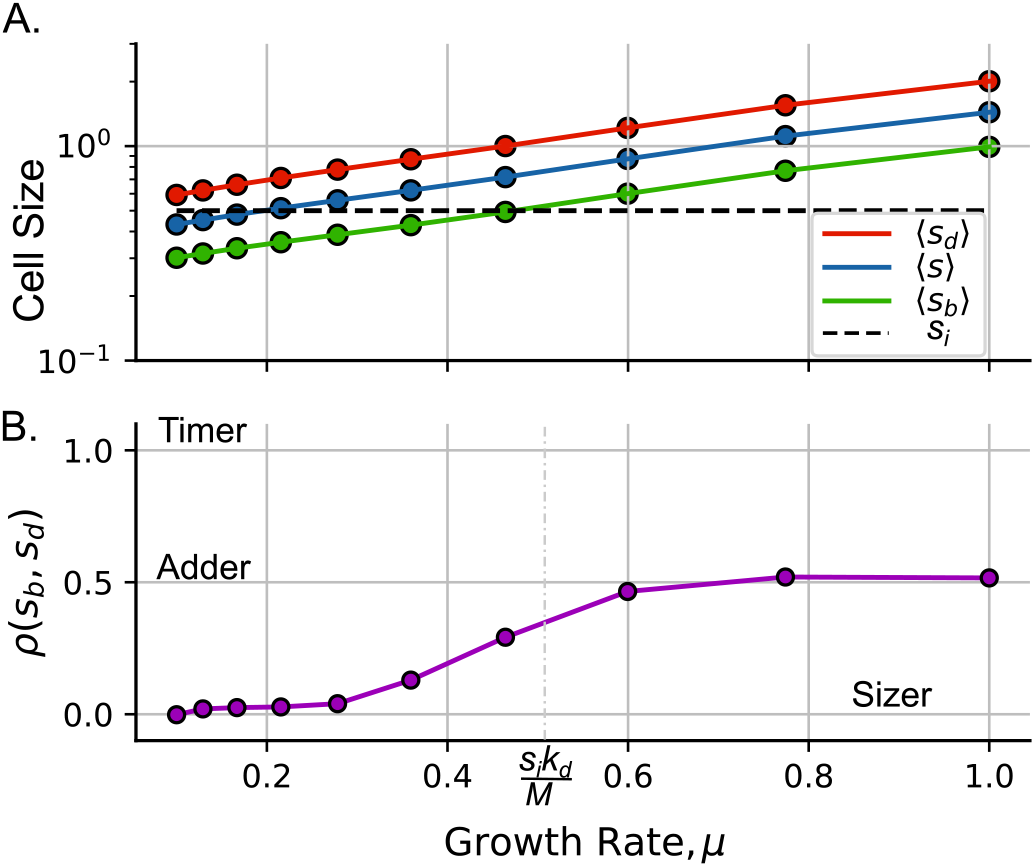
Size regulation in different steady growth conditions. A. Mean size ⟨*s*⟩ (blue), mean size at division ⟨*s*_*d*_⟩ (red), and mean size at birth ⟨*s*_*b*_⟩ (green) as a function of the growth rate. B. Correlation coefficient *ρ*(*s*_*b*_, *s*_*d*_) as a function of the growth rate. (Parameters: *s*_*i*_ = 0.5, *k*_*d*_ = *M, M* = 10)

Fig. 2B shows how *ρ* is null valued (sizer) when the growth rate satisfies 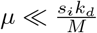. Then, for faster growth conditions 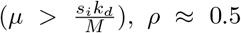, *ρ* ≈ 0.5. That is, the division strategy is an adder. The manner in which cells divide following the sizer strategy under slow growth conditions was previously found in experiments (Wallden et al., 2016). Although the minimum size model predicts the sizer strategy in slow growth, it is worth mentioning that there are alternative models that may explain this phenomenon (Nieto et al., 2020a; Si et al., 2019).

In this section, we study the division strategy under steady growth conditions. The next section will present how the division strategy changes when the growth rate is not steady but presents a dynamics similar to the one observed in the nutrient depletion by population growth.

### 2.2 Size regulation along the growth curve

In this section, we model experiments similar to those performed by (Bakshi et al., 2021). This experiment consists of trapping cells in a microfluidic device and exposing them to the depletion of nutrients present in population growth. This means that we are not interested in the population dynamics, but only in the effect of the growth rate changes in the cell size and division dynamics. We first present a model to obtain the dynamics of the nutrients and, therefore, the growth rate dynamics. After that, we will show the resulting regulation of cell size under these conditions of depletion of nutrients.

Consider that the biomass of the population 0 *< B <* 1 (in convenient units) increases exponentially at a rate *µ*. As a particular case of the model proposed by (Nev et al., 2021), we will consider that *µ* is proportional to the concentration of nutrients 0 *< N <* 1 (also in convenient units). If *N* is also depleted at a rate proportional to biomass, we propose the following system:

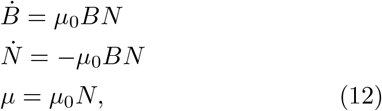

with *µ*_0_ a constant. The solutions for *B, N*, and *µ* follow:

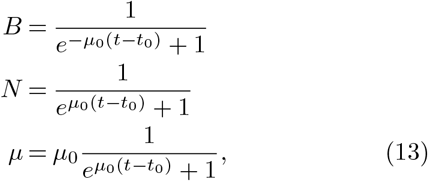

with 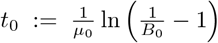 and 0 *< B*_0_ *<* 1 being the biomass at time *t* = 0.

Solutions condensed in (13) can be observed in Fig. 3. For example, Fig. 3A shows how *B* increases over time. On the other hand, the downward trajectory of *µ* is presented in Fig. 3B. Fig. 3A also shows the definition of exponential and stationary phases.

**Fig. 3.**
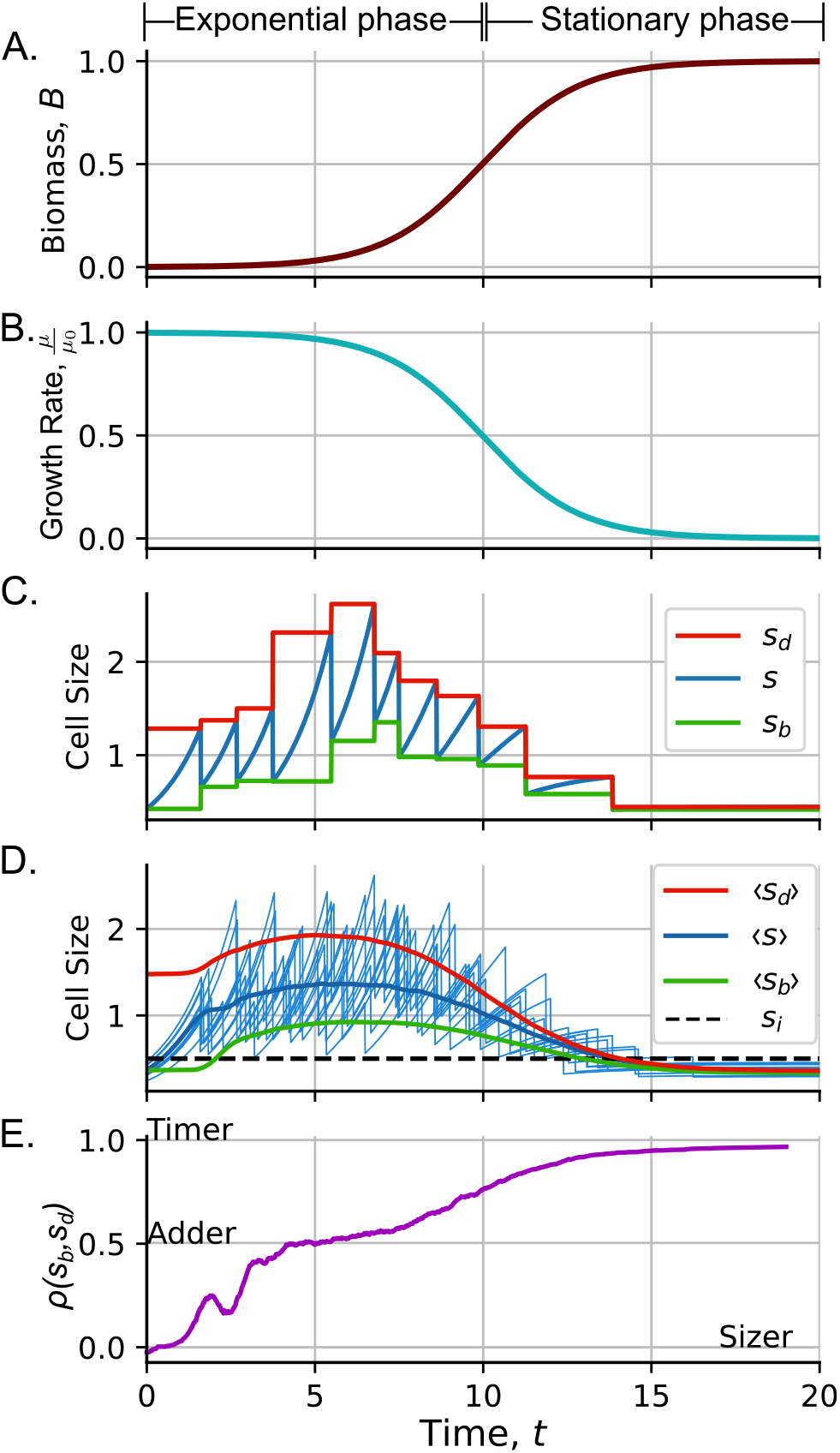
Dynamics of the cell variables during nutrient depletion by population growth. A. The biomass as a function of time. B. Dynamics of the growth rate. C. Cell size trajectory (blue), size at birth *s*_*b*_ (green) and the size at division *s*_*d*_ (red) for a single cell. D. Dynamics of the mean size ⟨*s*⟩ (blue) and some cell size trajectories (light blue on the background) as a function of time. Mean size at division ⟨*s*_*d*_⟩ (red) and mean size at birth ⟨*s*_*b*_⟩ along the time. The black dashed line represent the minimum size *s*_*i*_. E. Dynamics of the size strategy along the growth curve. 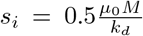, *M* = 10, *k*_*d*_ = 10 ln(2), *µ*_0_ = ln(2), *B*_0_ = 0.001, *N*_0_ = 1 − *B*_0_

To better illustrate the cell size dynamics, Fig. 3C shows a single cell size trajectory (blue). The initial conditions were fixed by making the cell face three times this growth curve before acquiring the data. To study the dynamics of the division strategy, we separated the time into cell-cycle intervals. Fig. 3C shows *s*_*d*_ for each cycle (red) and *s*_*b*_ (green).

Averaging five thousand cell size trajectories, we plot Fig. 3D presenting ⟨*s*⟩ (dark blue), ⟨*s*_*d*_⟩ (red), and ⟨*s*_*b*_⟩ (green) as a function of time. First, we can observe how entering the stationary phase makes the cell grow slower. Cells continue to divide, but since cell division stops only at the minimum size *s*_*i*_, they will have sizes around *s*_*i*_ when reaching the stationary phase.

The division strategy profile also shows a wide range of dynamics that is very similar to those found in experiments (Bakshi et al., 2021). When leaving the stationary phase, the cells show a sizer division (*ρ* ≈ 0). An intuitive explanation is that, at the end of the stationary phase, most cells are smaller than *s*_*i*_. When fresh medium is added to start the exponential phase, cell division begins only after meeting this minimum size. Once the division starts at *s*_*i*_ and not at *s*_*b*_, considering that the added size Δ is constant relative to *s*_*i*_, and since *s*_*i*_ is independent of *s*_*b*_, we expect *s*_*d*_ to be independent of *s*_*b*_.

After approximately three cycles, cells are definitely in balanced growth reaching the adder strategy *ρ* ≈ 0.5.This balanced growth continues until the cells decrease their growth rate. Thus, the division begins to behave like a timer *ρ* ≈ 1. Cells keep this strategy until the end of the stationary phase. A possible explanation for the timer division is that cells grow very slowly; that is, *s* is also approximately constant. As a result, the division rate *k*_*d*_*s* is also almost constant and cells divide, on average, after a given time (Sauls et al., 2016).

In this section, we observe how the model for minimum division size predicts the decrease of size entering the stationary phase and the transition from sizer to adder in the exponential phase and from adder to timer after entering the stationary phase. In the next section, we will present the size dynamics of cells growing in an arbitrary dynamic environment.

### 2.3 Size regulation in and arbitrary dynamical environment

To better understand the effects of changes on the growth rate, we propose a scenario in which *µ* varies dynamically. As Fig. 4A shows, we consider, at the beginning of the experiment, that cells are growing and dividing in a balanced way. The growth rate is such as 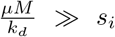. This environment is followed by a slow growth condition where 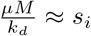, then the initial growth condition is again followed by a faster growth condition and the return to the initial state.

**Fig. 4.**
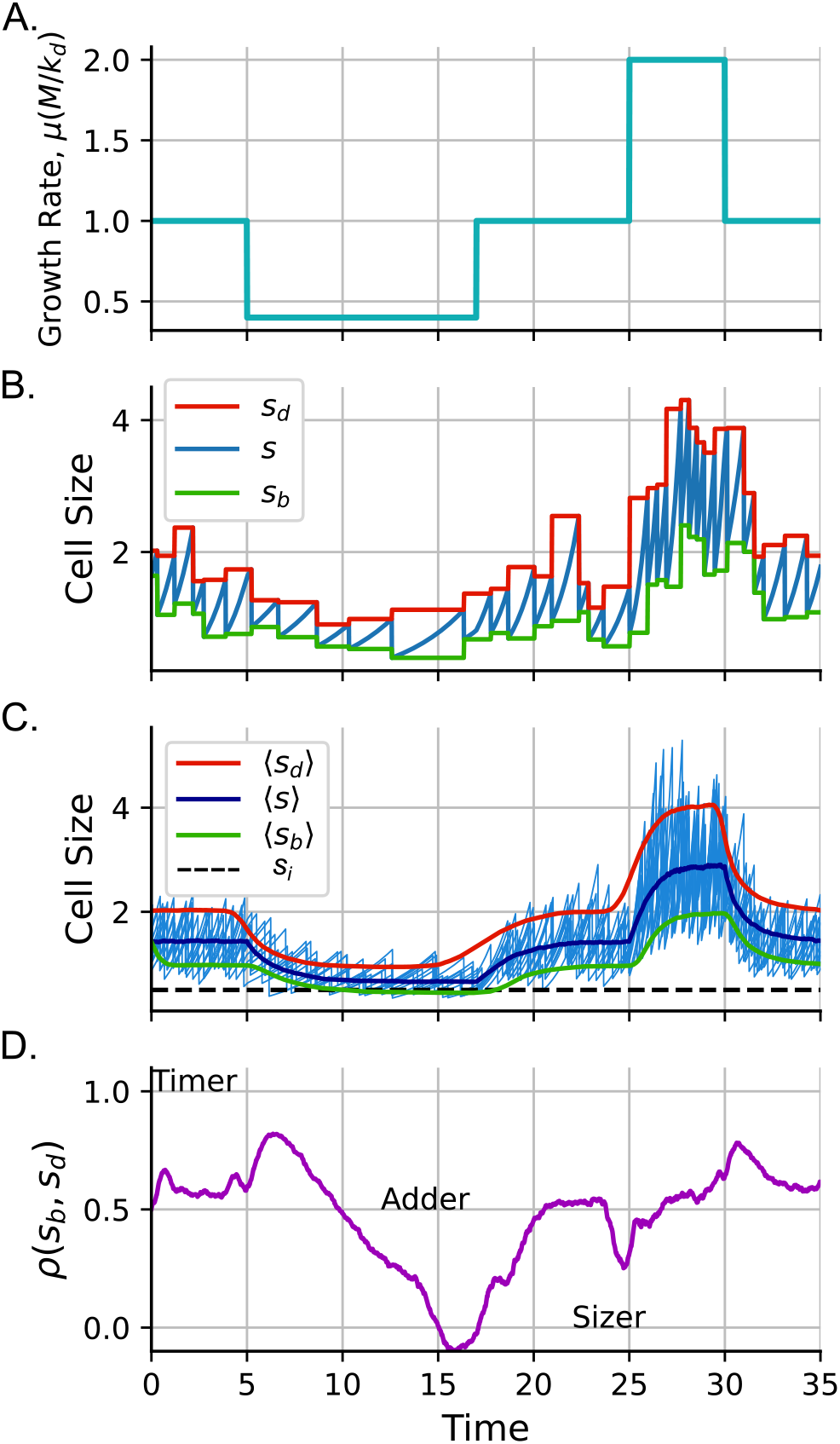
Dynamics of the cell variables along the growth curve. A. Arbitrary growth rate dynamics as a function of time. B. Mean size trajectory (blue), size at birth *s*_*b*_ (green) and the size at division *s*_*d*_ (red). C. Dynamics of the mean size ⟨*s*⟩ (blue) and some cell size trajectories (light blue on the background) as a function of time. Mean size at division ⟨*s*_*d*_⟩ (red) and mean size at birth ⟨*s*_*b*_⟩ along the time. The black dashed line represent the minimum size *s*_*i*_. D. Dynamics of the size strategy for this arbitrary dynamic environment. 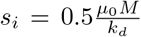, *M* = 10, *k*_*d*_ = 10 ln(2), *µ*_0_ = ln(2)

Fig. 4B shows the size trajectory of a single cell under these environment dynamics. Observing the dynamics of the mean size values (Fig. 4C), we observe how cells become smaller under slow growth conditions and recover their initial dimensions after growing under the initial conditions again. Finally, they become larger with faster growth, recovering their initial size after returning to the same growth rate from which they started.

The dynamics of the division strategy show an interesting behavior. They start from the adder strategy *ρ* ≈ 0.5 in balanced growth. While *µ* decreases, as in the growth curve, the division strategy moves closer to the timer. However, if we keep these slow conditions long enough, this correlation tends to the value observed in steady growth (Fig. 2B). When we suddenly transition from this slow growth condition to a faster one, we see how this correlation decreases for some small time interval. If these fast growth conditions are maintained, *ρ* relaxes again to the steady values shown in (Fig. 2B).

This behavior shows how sudden changes in environmental conditions can alter the transient dynamics of variables associated with the regulation of the splitting time. Additional comparisons with experiments can reveal the level of validity of this simple model.

## 3. DISCUSSION

The model we propose aims to describe the cell size dynamics. A direct comparison of these results with actual data must include additional variables. For example, in our model, we consider that there is no random variability in the growth rate as other models (Kiviet et al., 2014; Vadia and Levin, 2015; Nieto et al., 2022a). Another simplification of the model is that cell division starts exactly after reaching *s*_*i*_. Previous studies propose that a realistic model can consider the division based on the size following a Hill function with finite exponent (Nieto et al., 2020b).

Cell variables may show dynamic changes in their amplitude of random fluctuations. For example, near the stationary phase, gene expression has been reported to be more noisy due to activation of stress response mechanisms (Navarro Llorens et al., 2010; Bakshi et al., 2021). Therefore, a general model considering size variability must include dynamic changes in noise in variables such as the growth rate or the splitting position due to this stress response.

In dynamic environments, cell physiology can be altered in more complex subtle ways. For example, some studies show that during environmental changes, the synthesis of the cell wall and bulk proteins has different dynamics (Harris and Theriot, 2018; Sekar et al., 2018; Büke et al., 2022). These delays lead to changes in the shape of the cells that are not considered in this model.

An intriguing property of our model is that the division constant *k*_*d*_ is independent of the growth condition. One possible reason behind it is that the cell may consider splitting as a priority with properties not changing with the growth condition. This can be associated with the robustness of the replication and septal ring formation processes under different growth conditions (Shimaya et al., 2021; Witz et al., 2019; Sanders et al., 2022).

The interpretation of the mechanisms that govern size regulation is still under debate. The classification of division strategies in Timer, Adder, and Sizer is primarily pedagogical. Its definition is related to exponential growth and stable conditions. For example, if growth is linear, all these division strategies are equivalent (Totis et al., 2020). Therefore, although it is a useful quantitative tool for parameter inference (Munsky et al., 2018), the interpretation of division strategies at a nonexponential growth rate is sometimes not intuitive.

Among other limitations of the model, we have to note that we compare the results of our model with observations of cells that grow and divide without considering cell proliferation (Nieto et al., 2022b). Division affects not only size regulation, but also population dynamics, which may be the subject of future research.

Finally, understanding cell regulation in these complex dynamic environments can help us infer the mechanisms that govern not only cell size regulation but also other physiological phenomena associated with size, such as gene expression, stress response, and proliferation mechanisms (Zheng et al., 2016; Jun et al., 2018).

## 4. CONCLUSIONS

In this article, we study the dynamics of size regulation for cells that grow and divide in complex and dynamic environments. First, we propose a model of cell division that considers this event as a stochastic process with a rate proportional to the cell size. Its proportionality constant does not depend on the growth conditions and is null-valued if the cell size is smaller than a minimum size. From this model, we predict that the mean cell size is exponentially dependent on the growth rate. Also, we predict, as observed in experiments, that fast-growing cells follow the adder strategy, while slow-growing cells follow the sizer strategy.

We also study how the size dynamics and the division strategy change across the growth curve. Our model predicts that the first divisions after the exit of the stationary phase show a sizer strategy. In balanced growth, cell splitting follows the adder, and entering the stationary phase shows timer properties. These predictions also agree with the experimental results.

Finally, we study cell size dynamics and division strategy in an arbitrary fluctuating environment. As a principal conclusion, the model predicts that cell size decreases with slow growth and increases under fast growth conditions. If these conditions are persistent enough, the division strategy can reach the steady conditions observed in the first part of the article. The division strategy is closer to a timer during the transition from fast to slower growth conditions. The division strategy is closer to a sizer for a brief period when switching to a faster growth condition.

